# Comparison Between UMAP and t-SNE for Multiplex-Immunofluorescence Derived Single-Cell Data from Tissue Sections

**DOI:** 10.1101/549659

**Authors:** Duoduo Wu, Joe Yeong Poh Sheng, Grace Tan Su-En, Marion Chevrier, Josh Loh Jie Hua, Tony Lim Kiat Hon, Jinmiao Chen

## Abstract

Using human hepatocellular carcinoma (HCC) tissue samples stained with seven immune markers including one nuclear counterstain, we compared and evaluated the use of a new dimensionality reduction technique called Uniform Manifold Approximation and Projection (UMAP), as an alternative to t-Distributed Stochastic Neighbor Embedding (t-SNE) in analysing multiplex-immunofluorescence (mIF) derived single-cell data. We adopted an unsupervised clustering algorithm called FlowSOM to identify eight major cell types present in human HCC tissues. UMAP and t-SNE were ran independently on the dataset to qualitatively compare the distribution of clustered cell types in both reduced dimensions. Our comparison shows that UMAP is superior in runtime. Both techniques provide similar arrangements of cell clusters, with the key difference being UMAP’s extensive characteristic branching. Most interestingly, UMAP’s branching was able to highlight biological lineages, especially in identifying potential hybrid tumour cells (HTC). Survival analysis shows patients with higher proportion of HTC have a worse prognosis (*p*-value = 0.019). We conclude that both techniques are similar in their visualisation capabilities, but UMAP has a clear advantage over t-SNE in runtime, making it highly plausible to employ UMAP as an alternative to t-SNE in mIF data analysis.

## Introduction

Since the advent of immunotherapy, analyses of the tissue immune microenvironment have become a critical bridge between cancer research and patient care. Tumour infiltrating lymphocytes (TILs) are highly heterogeneous at the single-cell level; consequently, single-cell analytical techniques such as flow/mass cytometry and single-cell RNA-sequencing (scRNA-seq) are essential to study the immune microenvironment. New single-cell flow cytometry techniques, such as the BD FACSymphony(tm) ^1^ system has greatly increased the number of immunological parameters that can be measured per cell, from 10 to 27. Mass cytometry can simultaneously measure more than 40 markers per cell while single-cell RNA-seq reports the entire transcriptome of individual cells. Despite these advances, location information for individual cells is lost. By contrast, immunohistochemical techniques capture spatial information for individual cells, but only a few markers can be measured at one time. Immunohistochemical technologies are also advancing, with the latest multiplex-immunofluorescence (mIF) system able to measure seven markers, and Imaging Mass Cytometry systems (incorporating high-parameter CyTOF® technology with imaging capability) able to process up to 37 biomarkers simultaneously.

Dimensionality reduction is critical for all of these single-cell technologies to reduce the number of variables (dimensions) under consideration in the samples. Dimensionality reduction techniques include Principal Component Analysis (PCA), ISOMAP^2^, diffusion map ^3^ and t-distributed stochastic neighbour embedding (t-SNE) ^4^, which have been established to visualize single-cell data. Among these methods, t-SNE is relatively well-studied and the most commonly used. In 2018, McInnes and Healy published a new dimensionality reduction method known as uniform manifold approximation and projection (UMAP) ^5^. UMAP is effective for mass cytometry and single-cell RNA-seq data^6^, and the researchers state that it has a shorter running time and better preservation of the global structure of the data than tSNE ^5^.

Here, we compared the utility of UMAP with tSNE on mIF data. To evaluate the biological relevance of UMAP as a dimensionality reduction tool for mIF data, we adopted the FlowSOM unsupervised clustering method ^7^ to identify eight major cell types present in human hepatocellular carcinoma (HCC) tissues. We used UMAP and t-SNE independently on the same mIF dataset to qualitatively compare the distributions of the FlowSOM-identified cell types produced by both dimensionality reduction tools.

Lastly, we built an integrated R package called “Harmony” for an automated analysis pipeline of mIF data. This pipeline includes data pre-processing, dimensionality reduction using t-SNE and UMAP, clustering analysis and visualization of the spatial distribution of the various cell types.

## Methods & Materials

### Patients and tumours

A total of 165 archived formalin-fixed, paraffin-embedded (FFPE) HCC specimens from patients diagnosed between May 1997 and July 2007 at the Department of Anatomical Pathology, Division of Pathology, Singapore General Hospital, were analysed (Figure 1). All samples were obtained before patients underwent chemotherapy or radiotherapy (Table 1), and clinicopathological parameters, including patient age, tumour size, histologic growth pattern, grade and subtype, lymphovascular invasion and lymph node status were collected. The patient age ranged from 15-88 years (median, 64 years) while length of follow-up ranged from 1-159 months (mean, 57 months; median, 44 months); recurrence and death occurred in 32 (19%) and 58 (35%) of these patients, respectively. Tumours were typed, staged and graded according to the World Health Organization, American Society of Clinical Oncology-College of American Pathologists (ASCO-CAP) guidelines ^8^. Ishak Fibrosis Score ^9, 10^, which was documented in the pathological diagnostic reports, was used to evaluate the fibrosis status of the non-neoplastic liver. More information regarding patient demographics is displayed in Table 1. The Centralized Institutional Review Board of SingHealth provided ethical approval for the use of patient materials in this study (CIRB ref: 2009/907/B).

**Table 1:**
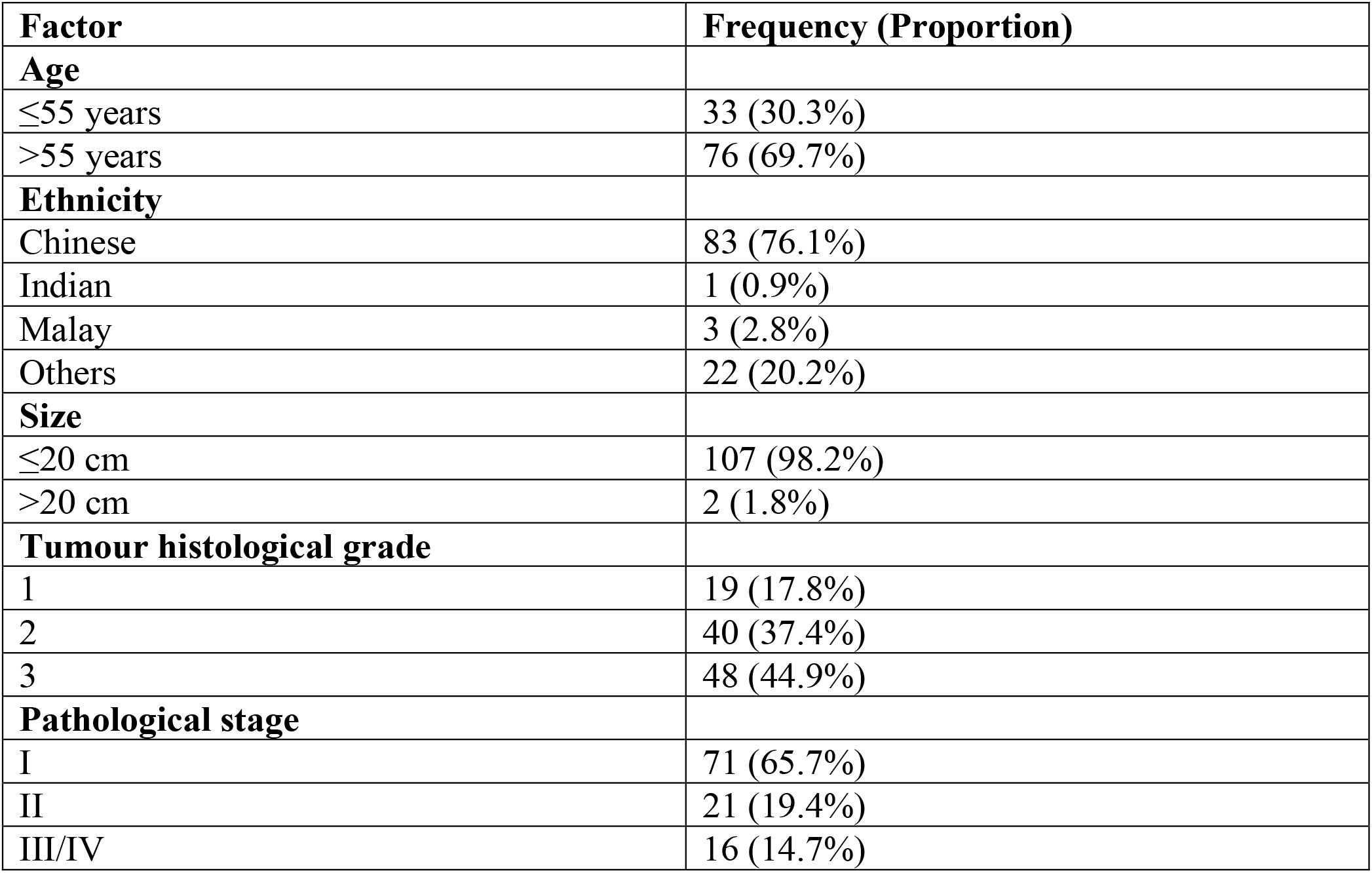
Summary of patient demographics

**Figure 1:**
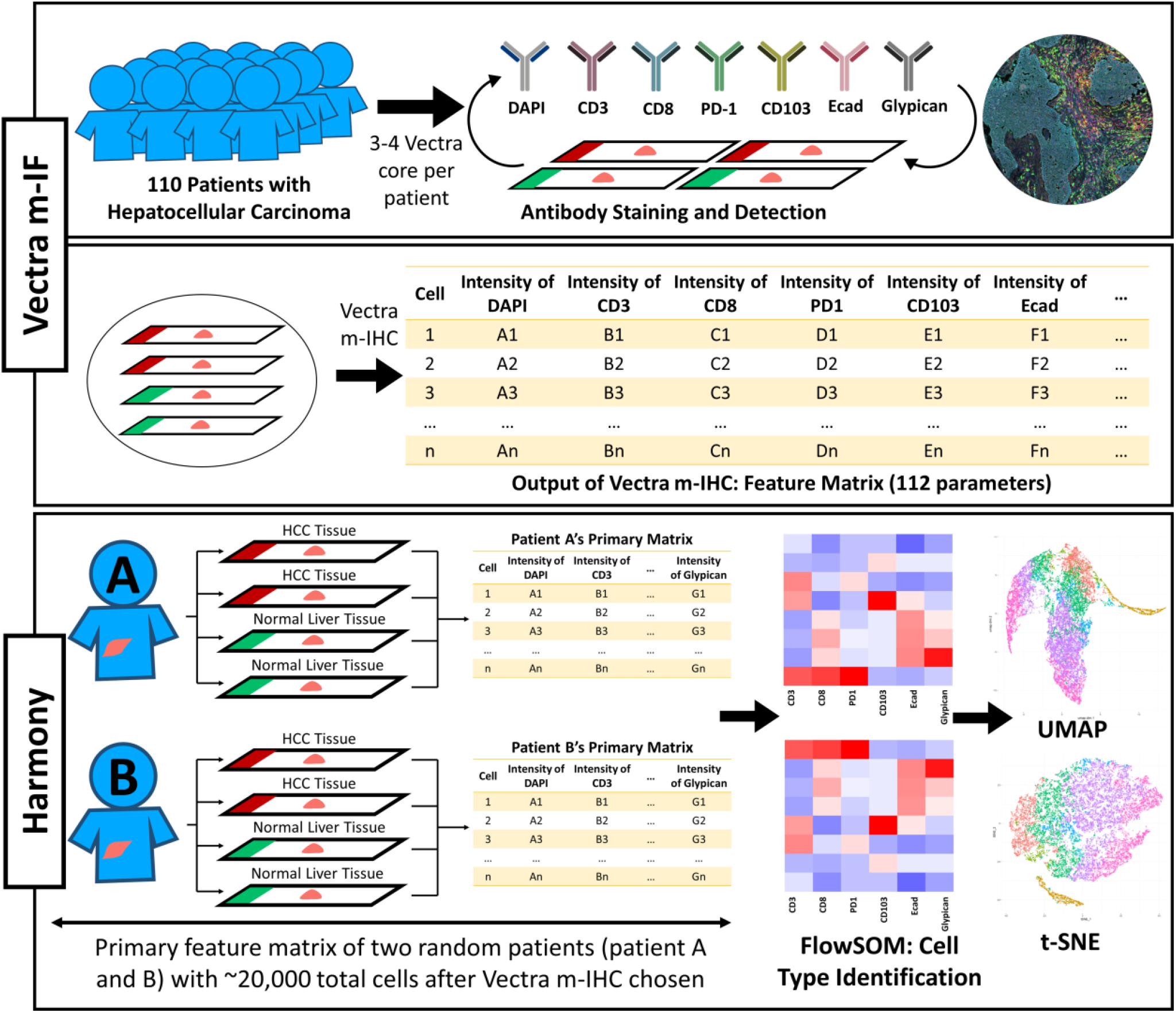
Overview of methodology.

### Tissue microarray (TMA) construction

Tumour (where >50% of the sample area was tumour tissue) and adjacent normal liver regions for TMA construction were selected by pathological assessment. For each sample, two or three representative tumour cores of 1 mm diameter were transferred from donor FFPE tissue blocks to recipient TMA blocks using an MTA-1 Manual Tissue Arrayer (Beecher Instruments, Inc., Sun Prairie, WI, USA). TMAs were constructed as previously described ^11^.

### Quantitative Multiplex immunofluorescence (QmIF)

Quantitative multiplex immunofluorescence (QmIF) was performed using an Opal Multiplex Immunohistochemistry (IHC) kit (PerkinElmer, Inc., Waltham, MA, USA) as previously described ^12-22^, on FFPE tissue sections processed according to the standard IHC protocol described above. The slides were first incubated with primary antibodies against CD3, CD8, CD103, Glypican, Ecadherin and PD-1 and the nuclei were counterstained with DAPI (Table 2), before incubation with fluorophore-conjugated tyramide signal amplification buffer (PerkinElmer, Inc., Waltham, MA, USA). Images were captured under a Vectra 3 pathology imaging system microscope (PerkinElmer, Inc.) and analysed using inForm software (version 2.4.1; PerkinElmer, Inc.) ^23-25^. mIF returned a total of 112 measurements for each cell in the HCC samples, based on the mean, minimum, maximum and standard deviation of the seven markers in the cell membrane, cytoplasm, nucleus and entire cell (refer to supplementary Table 1).

**Table 2:** Summary of antibody markers used to stain the biopsies

| Antibody | Clone | Dilution | Source | Labelling Pattern |
| --- | --- | --- | --- | --- |
| CD3 | Polyclonal | 1:200 | Dako A0452 | membrane |
| CD8 | 4B11 | 1:100 | Leica Biosystems CD8-4B11-L-CE | membrane |
| CD103 | EPR4166(2) | 1:800 | Abcam (AB129202) | cytoplasmic, membrane |
| Glypican | 1G12 | 1:800 | Cell Marque 216M-96 | cytoplasmic, membrane |
| Ecadherin | NCH-38 | 1:30 | Dako M3612 | membrane |
| PD-1 | NAT105 | 1:100 | Cell Marque 315M-96 | cytoplasmic, membrane |

### FlowSOM Unsupervised Clustering

Two samples containing ∼20,000 cells were randomly chosen for comparison (patient A and B). These data were inputted into “Harmony” for the automated analytical pipeline. “Harmony” will first identify the cell types present in the two samples using FlowSOM, which uses an unsupervised clustering method based on self-organising maps (SOM) ^7^ and is typically used to analyse flow and/or mass cytometry data. FlowSOM was applied to raw untransformed data extracted from IF images using inForm software. The average marker expression in each FlowSOM cluster was visualized as a heatmap, from which an experienced pathologist assigned each FlowSOM cluster a biologically meaningful cell type (Figure 2). The annotated cell types were then overlaid on the t-SNE or UMAP reduced dimensions.

**Figure 2:**
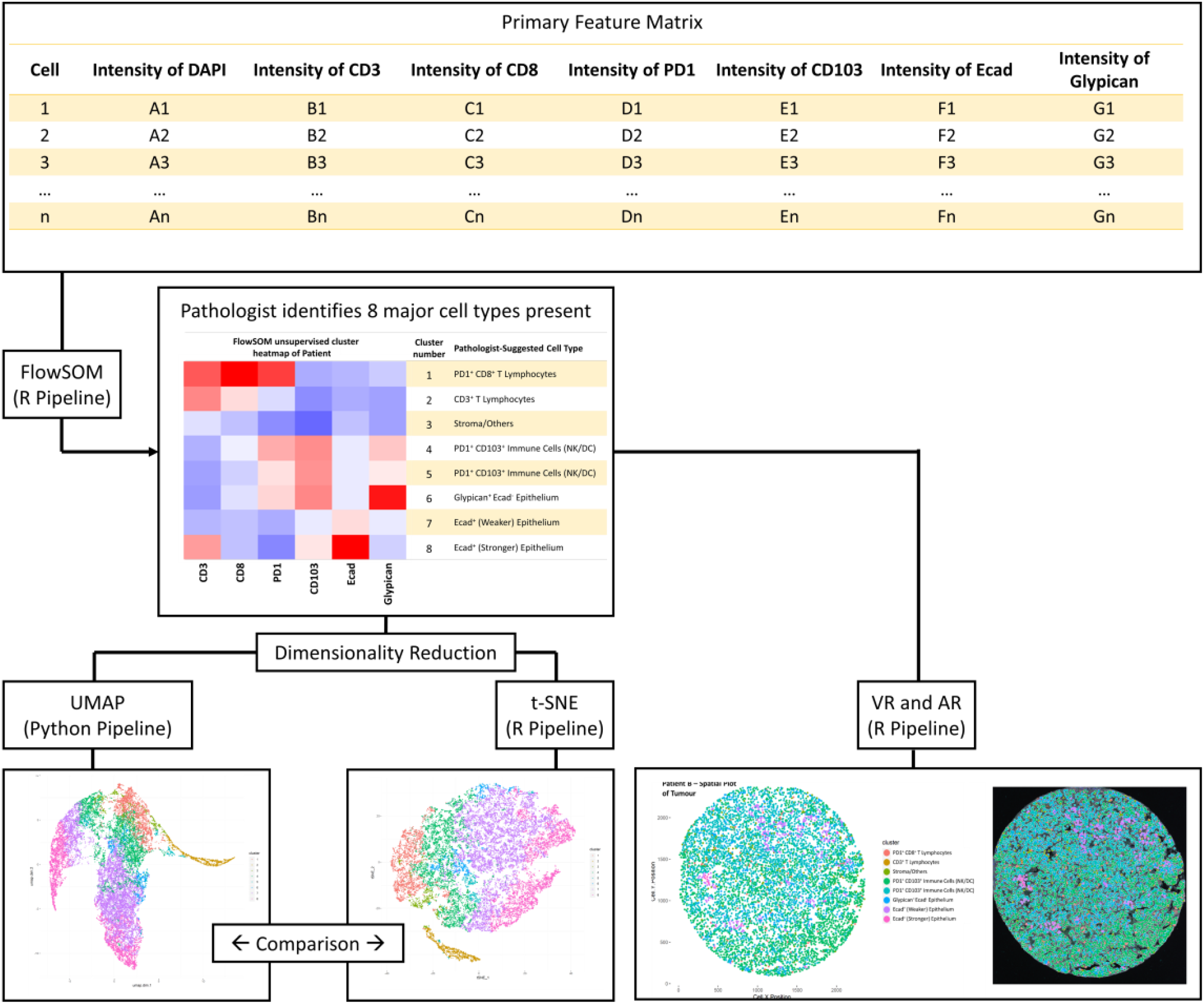
Analytical pipeline for “Harmony”.

### Dimensionality Reduction

UMAP and t-SNE were run independently using “Harmony” on the same dataset to qualitatively compare the distribution of clustered cell types in both dimensionality reduction tools. t-SNE was run using Rtsne, allowing it to be integrated into one seamless pipeline with FlowSOM. t-SNE was run without prior principal component analysis (pca = FALSE) due to the low number of biomarkers used (n= 7 biomarkers), and the perplexity levels were varied in multiples of 10 to identify the best visualisation. All remaining parameters were run using the default options. UMAP was run using a python script (https://github.com/lmcinnes/umap) but was integrated as part of “Harmony” R Package using the “Reticulate” library in R. Minimum distance of 0.3 was set for UMAP. The number of nearest-neighbours parameter, a parameter in UMAP equivalent to the perplexity parameter in t-SNE, varied from two to 20 until the best visualisation was obtained. All other parameters were run using the default options. All visualisations were plotted using R.

### Survival Analysis

Overall survival curves were estimated by the Kaplan-Meier method and compared using log-rank (Mantel-Cox) test. A p-value less than 0.05 was considered statistically significant. Patients who are still alive at the last follow-up were censored. Cox proportional hazards regression model was used to relate risk factors, considered simultaneously, to overall survival time. The parameter estimates represent the change in the expected log of the hazard rate relative to a one-unit change in the risk factor, holding all other risk factors constant. Tests of hypothesis were used to assess whether there are statistically significant associations between risk factors and time to event. A p-value less than 0.05 was considered statistically significant. The parameter estimates were generated using R 3.5.0 and are shown with their p-values, their associated hazard rates along with their 95% confidence intervals.

## Results

### Dimensionality reduction and unsupervised clustering with core features

We first conducted QmIF on tumour and normal hepatic tissue samples from two randomly selected patients (Figure 1). A sample of the patient’s tissue image is shown in Supplementary Figure 1. We then performed dimensionality reduction with both t-SNE and UMAP using only the mean intensities of the seven individual biomarkers in their respective cellular compartments for both patients (Table 3). Next, we applied FlowSOM to the original high-dimensional data, which categorised the cells into eight major clusters (Figure 2) and displayed the mean expression levels of the seven markers in each cluster in heatmaps (Figures 3a and 3b).

**Table 3:** List of measurements regarded as primary features

|  | Feature Measured |
| --- | --- |
| Primary Features (7 parameters) | Mean intensity of cytoplasm CD3 |
|  | Mean intensity of cytoplasm CD8 |
|  | Mean intensity of cytoplasm PD1 |
|  | Mean intensity of cytoplasm CD103 |
|  | Mean intensity of cell membrane Ecad |
|  | Mean intensity of cytoplasm Glypican |
|  | Mean intensity of nucleus DAPI |

**Figure 3:**
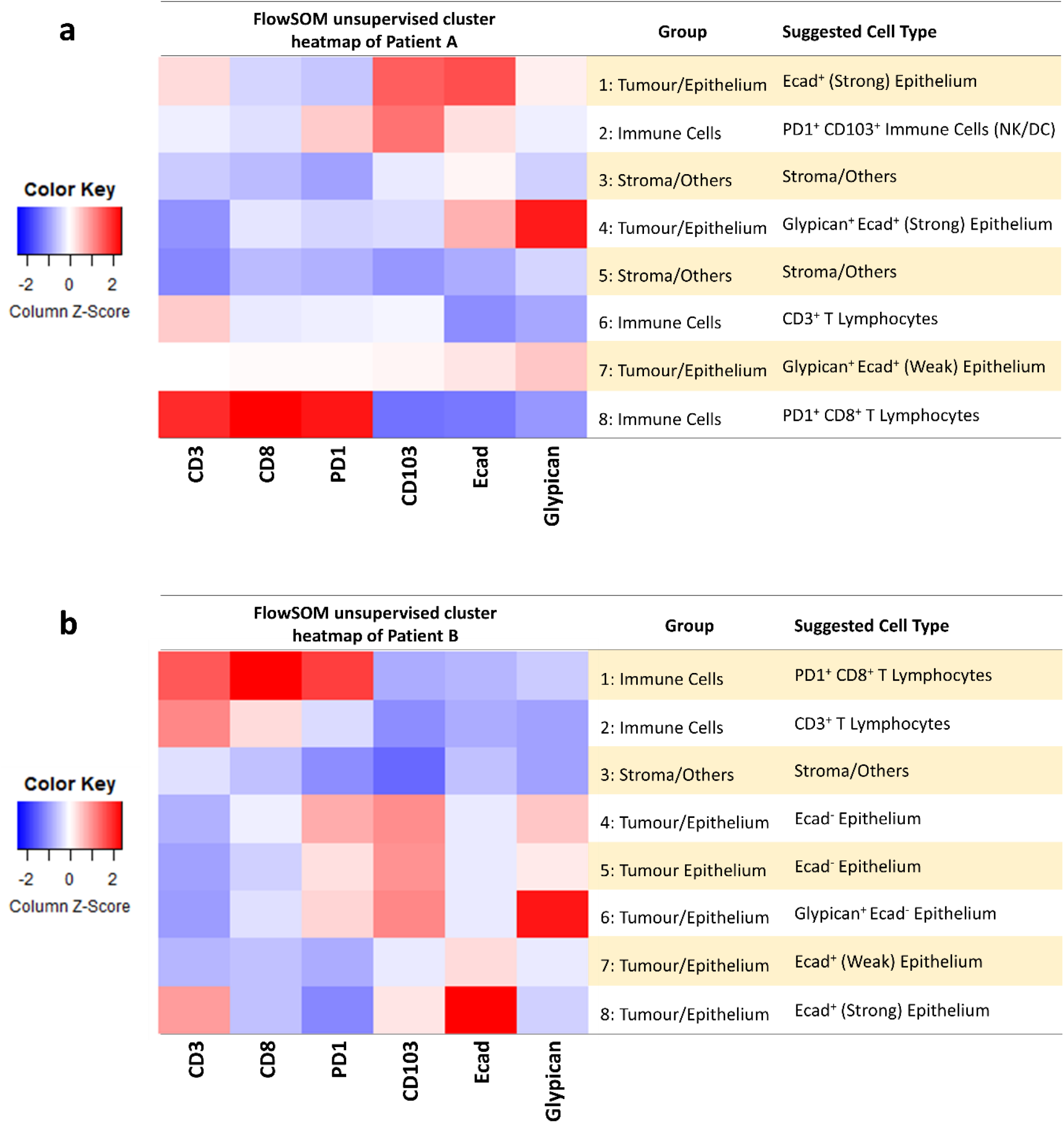
FlowSOM unsupervised cluster heatmap of a) Patient A and b) Patient B

The major cell types could be classified into three groups (Figures 3a and 3b): tumour/epithelial cells, immune cells and stromal/other cells. The tumour/epithelial cell clusters were either Glypican^+^ or Glypican^−^, and either Ecad^+^ or Ecad^−^. Ecad^+^ intensity could be further subclassified as either strong or weak. Based on this marker expression pattern, a Glypican^+^ epithelium most likely represents tumour cells, and the Ecad intensity indicates the characteristics of the epithelium (i.e. whether tumour cells have maintained or lost Ecad expression). For the immune compartment, the immune cells were either CD3^+^ (T lymphocytes), CD8^+^ (cytotoxic T lymphocytes) or CD103^+^ (macrophages or natural killer cells). Combining this information with the PD1 intensity revealed further cellular characteristics, such as the exhaustive status of the T lymphocytes. Clusters with low intensity marker expression across all seven markers were labelled as stromal/other cell types as none of the seven biomarkers were specific to these other cell types. The resulting dataset was subsequently dimensionally reduced using UMAP and t-SNE (Figure 2).

### UMAP runtime is significantly shorter than t-SNE at high cell numbers

We varied the t-SNE perplexity and the UMAP number of nearest-neighbours to study the effects of these parameters on runtime. We also varied the cell numbers in the dataset in intervals of a thousand from 1,000 to 19,000 to observe the change in time taken by both UMAP and t-SNE (Figure 4a). Here, we found that t-SNE ran faster than UMAP with low cell numbers (1,000 – 2,000 cells), whereas UMAP ran faster with higher cell numbers (>2,000 cells). We thus consider that running large datasets with UMAP will be more efficient than t-SNE in terms of runtime. Moreover, at t-SNE perplexity of 70 (a normal value of perplexity used to compute the visualisation), we observed that the UMAP runtime was an order of magnitude lower than t-SNE (Figure 4b).

**Figure 4:**
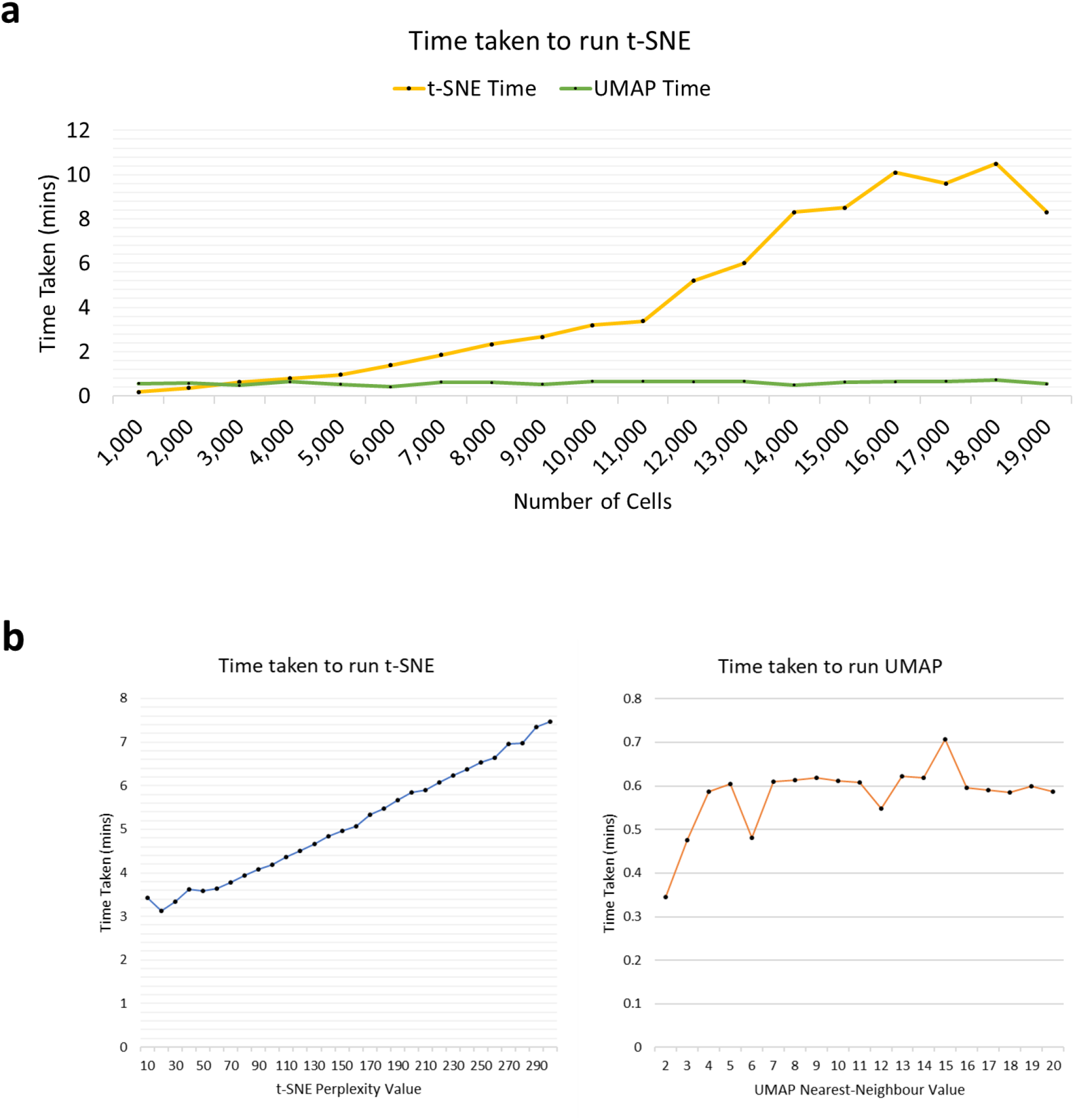
Assessment of time efficiency of t-SNE and UMAP. a) Comparison of runtime between UMAP and t-SNE with different numbers of total cells. b) Comparison of runtime of UMAP vs t-SNE given varying perplexity and number of nearest-neighbours values.

A new R pipeline, Fast Fourier Transform-accelerated Interpolation-based t-SNE (FIt-SNE), has been developed to reduce the runtime for t-SNE dimensionality reduction ^26^. When we applied FIt-SNE to our dataset with seven dimensions, however, we found that the original t-SNE algorithm was faster, taking only 6.2 mins compared to FIt-SNE taking 10.0 mins. Similarly, when we used the 112 dimensions from our dataset (Supplementary Table 1), FIt-SNE took 6.3 mins whereas the original t-SNE took only 4.5 mins. Although not effective on our data, FIt-SNE may be able to compute the visualisation with a reduced runtime when testing with higher dimensions (>112 parameters).

### t-SNE and UMAP produce similar cluster arrangements with comparable robustness and reproducibility

When visualizing the dimensionally reduced feature matrices, we observed that UMAP and t-SNE produced similar cluster arrangements (Figure 5). In particular, the major morphological clusters (clusters 1 – 9; Figure 5) produced by t-SNE and UMAP shared similar shapes and locations. In addition, we found the arrangements of cell types within the clusters were congruent between both visualisations. For example, t-SNE and UMAP visualisations of Patient B both produced a distinctive wedge-shaped cluster 5, with CD3^+^ T lymphocytes lying strictly adjacent to PD1^+^ CD8^+^ T lymphocytes and stromal cells, but the PD1^+^ CD8^+^ T lymphocytes not in contact with the stromal cells (Figure 5c and 5d). Additionally, both UMAP and t-SNE clustered tumour and normal hepatic cells from the two patients into different clusters (Figure 6). These data show that both methods can pick up differences in biomarker levels that permit clustering of tumour and normal cells to different groups. This is unlikely to be due to batch effect as this clear segregation of tumour cells from normal cells was evident for both patients.

**Figure 5:**
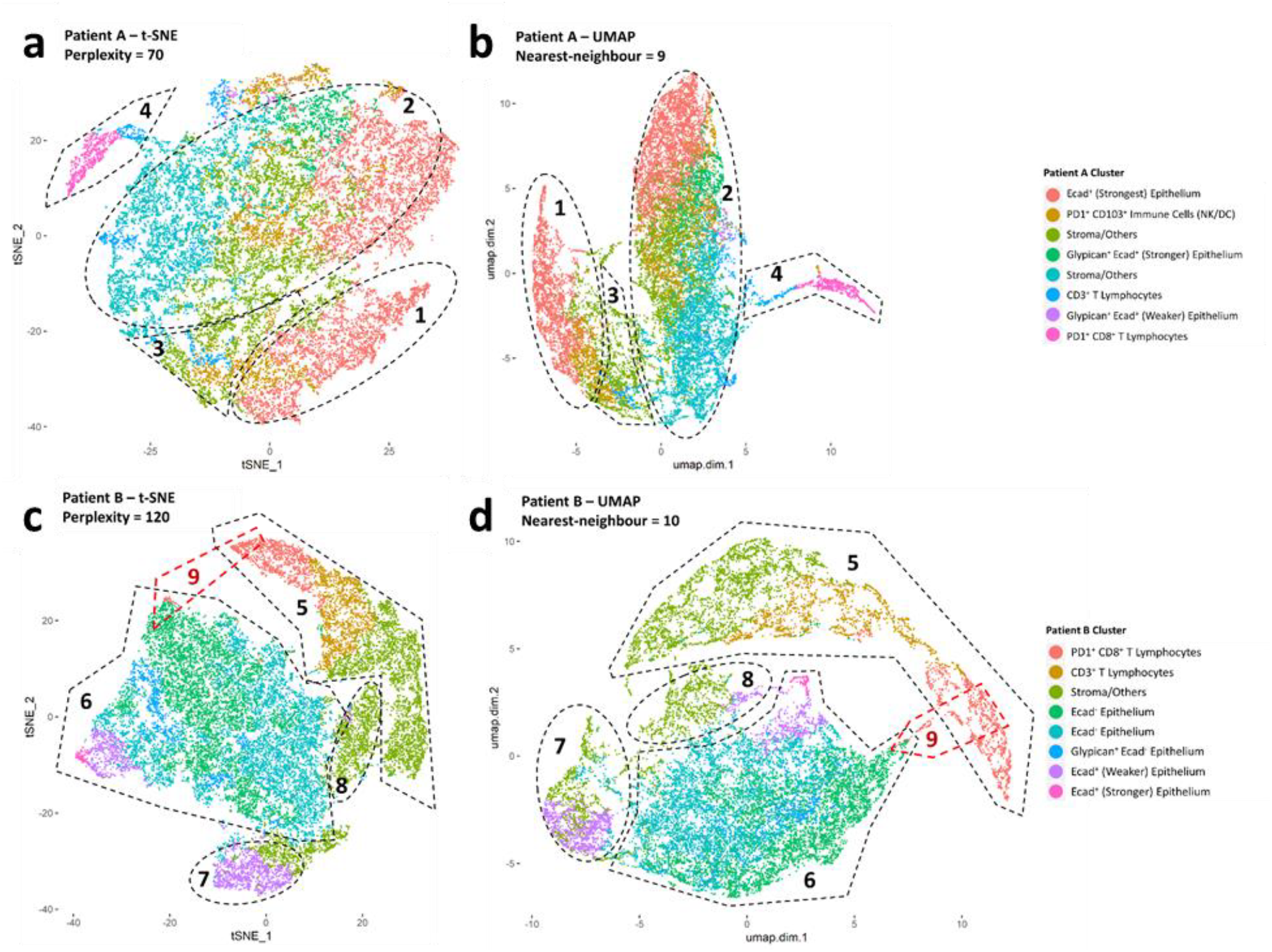
t-SNE and UMAP of patient A and patient B highlighting various cell types present. Number of nearest neighbours for UMAP and perplexity for t-SNE were varied until a good visualisation is obtained. a) t-SNE of patient A; b) UMAP of patient A; c) t-SNE of patient B; d) UMAP of patient B.

**Figure 6:**
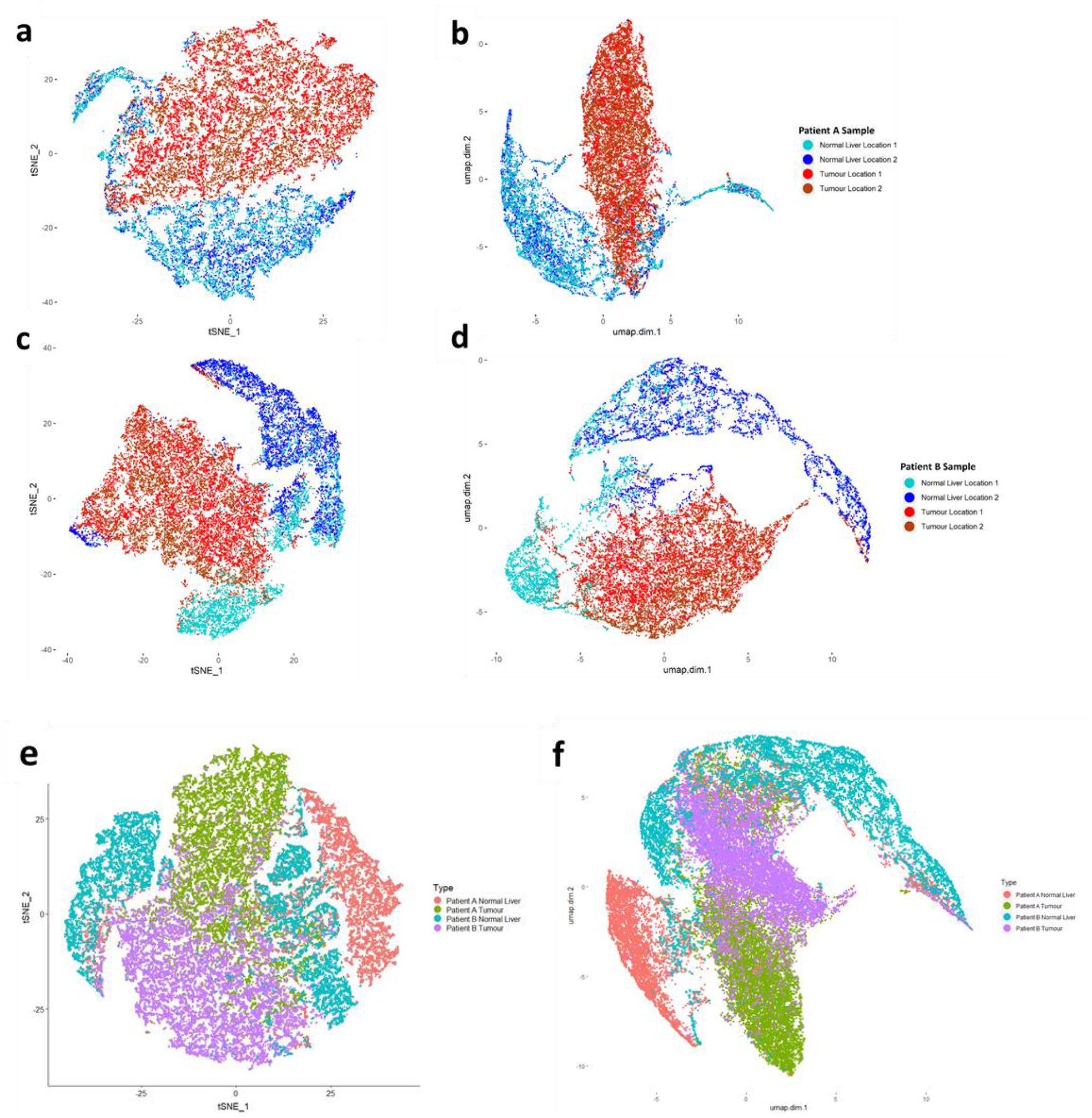
t-SNE and UMAP of patient A and patient B showing sources of various cells. a) t-SNE of patient A; b) UMAP of patient A; c) t-SNE of patient B; d) UMAP of patient B; e) combined t-SNE of patient A and B; f) combined UMAP of patient A and B.

We did identify morphological variations in the visualisations between the two techniques, with differences in the shape and point densities (Figure 5). Notably, the UMAP visualisations had signature branching characteristics, with continuous and “curvy” point plots. These point plots were more continuous than those produced by t-SNE, tending to form connections between clusters and cell types in the visualisations (Figure 5b and 5d). For example, UMAP preserved the cell lineage of the PD1^+^ CD8^+^ T cells in Patient B cluster 9 by forming a connection between cluster 5 and the Ecad^−^ epithelium cells in cluster 6; these connections were not produced by t-SNE. These cells in cluster 9 may either be immune cells that have lost their immune markers on cell surfaces, or tumour cells that have just started to express immune markers.

Furthermore, using additional secondary features (n=112), based on the mean, maximum, minimum and standard deviation of marker intensities measured at the nucleus, cytoplasm, cell membrane and entire cell created a t-SNE plot with virtually the same morphology as the UMAP visualisation (Figure 7).

**Figure 7:**
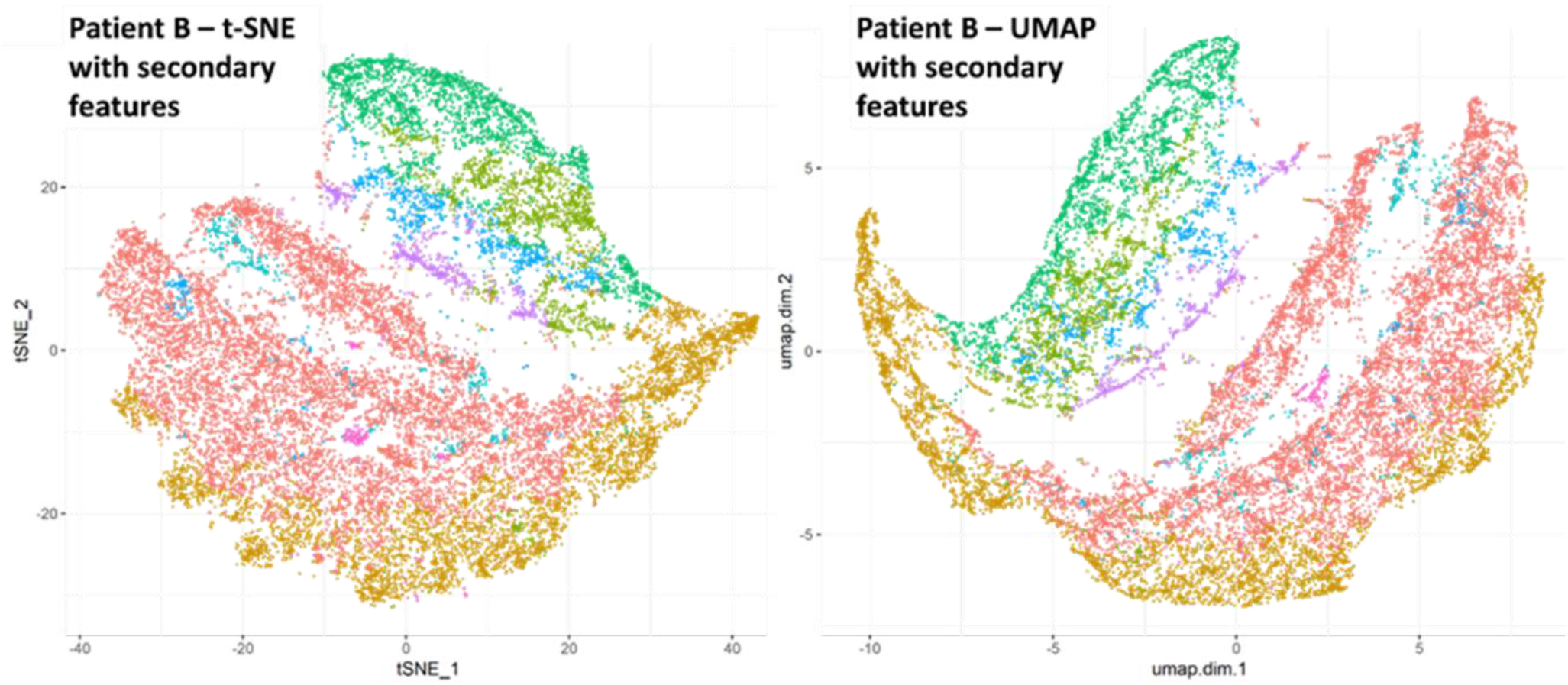
UMAP and t-SNE of patient B with 112 parameters.

We next randomly selected 90% of cells from Patient B to test the robustness and reproducibility of UMAP and t-SNE (Figures 8a and 8b). On a qualitative basis, both methods showed consistent clustering and cell-type arrangements. Most interestingly, the connection between the PD1^+^ CD8^+^ T cells and the Ecad^−^ epithelium cells was re-produced in each UMAP plot. However, it was also noted that relative physical locations of individual clusters varied across all the plots, especially the purple clusters.

**Figure 8:**
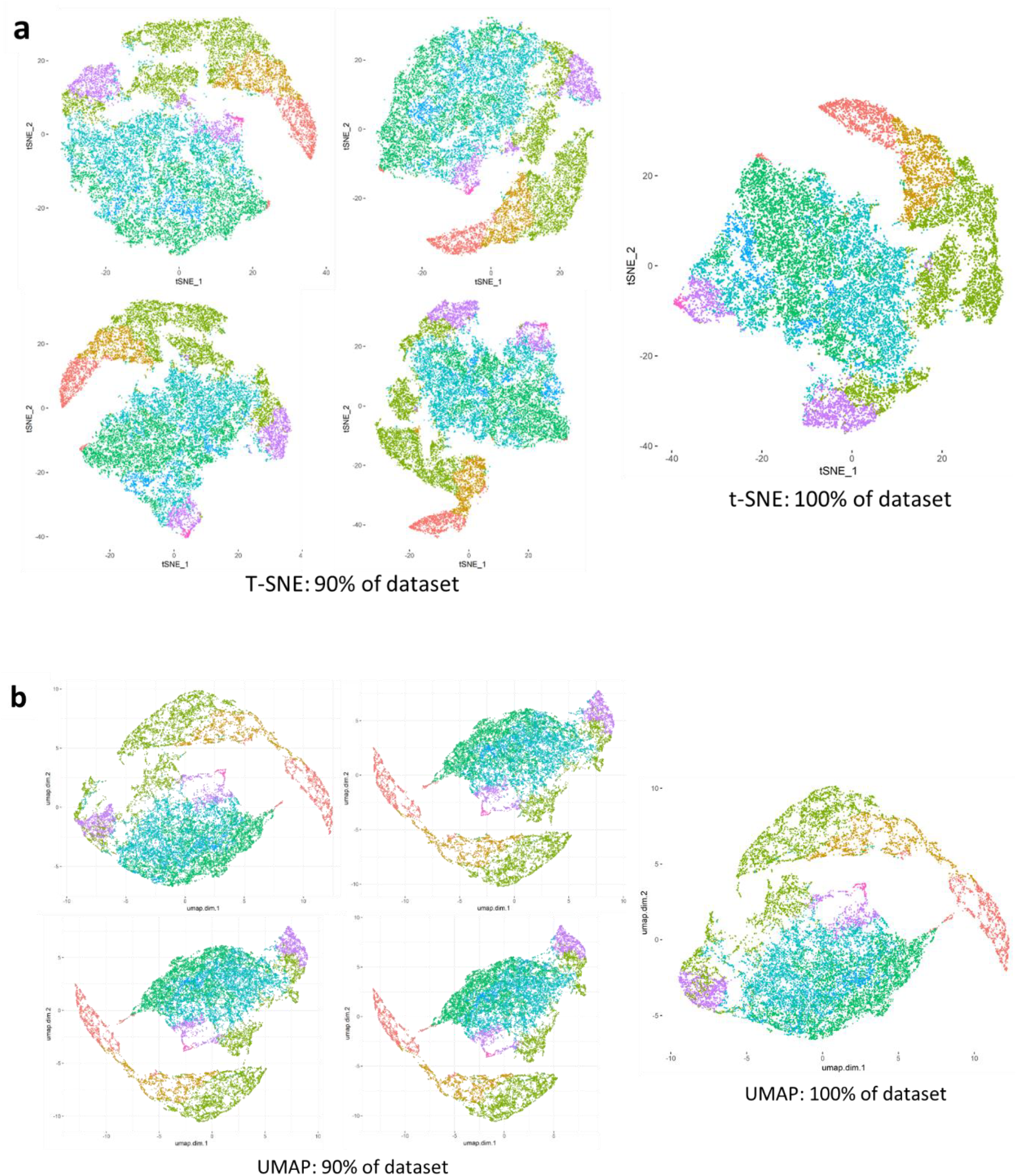
Assessment of robustness and reproducibility of t-SNE and UMAP. a) t-SNE of patient B’s dataset with 90% of cells randomly selected; b) UMAP of patient B’s dataset with 90% of cells randomly selected.

### Potential in identifying hybrid tumour cells and their association to patient prognosis

A recent study showed that tumour cells express immune markers to evade detection by the body’s immune system^28^. Our UMAP visualizations also detected cells with tumour-immune hybrid phenotypes as shown by the bridging between the PD1^+^ CD8^+^ T cells and the Ecad^−^ epithelium cells (cluster 9 of Figure 5d). If this is true, then UMAP should be superior to t-SNE when showing cell lineages. To identify the bridging cells shown in cluster 9, heatmaps of the various biomarkers were visualized on UMAP plots (Figure 9). It is observed that these cells are PD1^+^, CD8^+^, Glypican^+^ and CD103^+^ (Figure 9). These cells expressed both tumour and immune markers. Cut-off values provided by pathologists of 1.4, 20, 5.4, 5.0 and 4.0 were used for CD103^+^, CD8^+^, PD1^+^, Ecad^+^ and Glypican^+^ respectively. The percentages of these suspected hybrid tumour cells were calculated in all 119 biopsies in our dataset. Subsequently, Kaplan-Meyer survival curves were constructed, showing that biopsies with a higher proportion of these suspected hybrid cells had a poorer prognosis than those who had a lower proportion of these cells (*p*-value = 0.019) (Figure 10) by using a cut-off between “High” and “Low” proportion of hybrid cells equal to 1.7%. Among the 119 biopsies included and with a median follow-up of 73 months (4-160), 5-year overall survival was 39.5%, 95% CI = [22.3;70.0] in High hybrid cell proportion and 57.7%, 95% CI = [45.1;67.9] in Low hybrid cell proportion. Median OS was 42.7 months, 95% CI = [16.7; not reached] in High hybrid cell proportion and 117.1 months, 95% CI = [54.1; not reached] in Low hybrid values (Figure 10).

**Figure 9:**
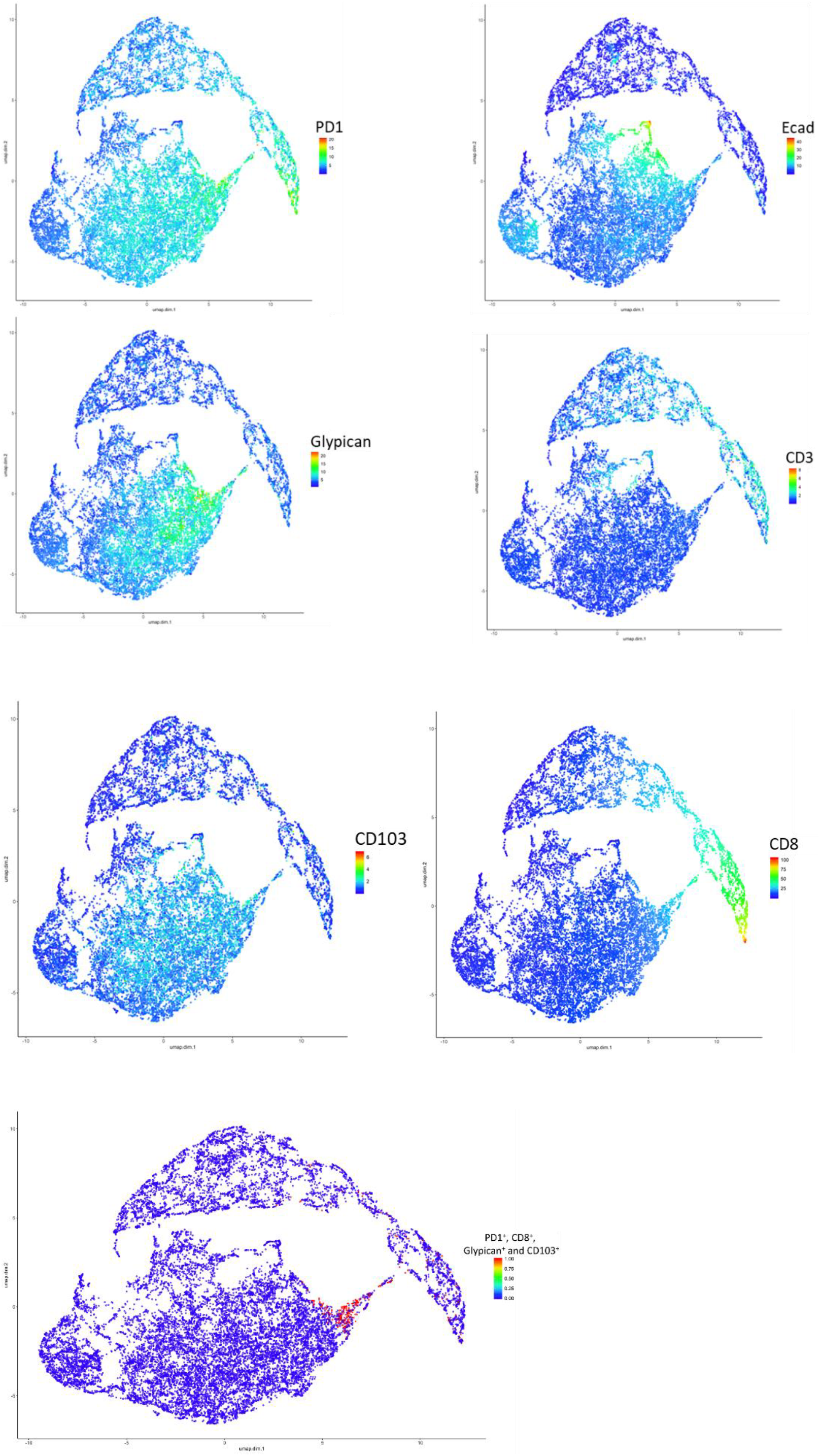
UMAP of patient B highlighting marker intensity in various locations of the UMAP plot. It was determined that the bridging cells are PD1^+^, CD8^+^, Glypican^+^ and CD103^+^.

**Figure 10:**
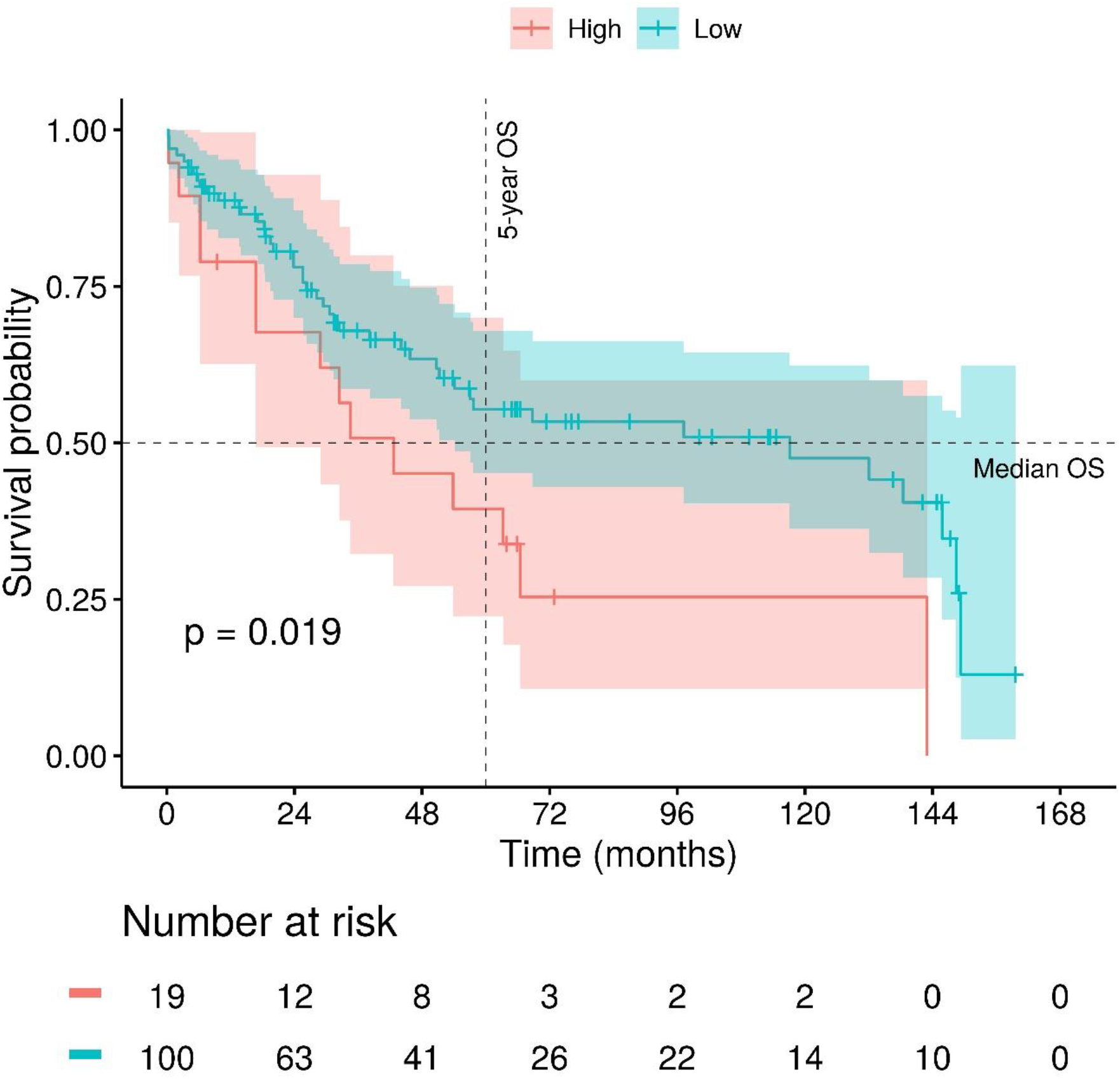
Kaplan-Meier survival curves showing overall survival in months with proportion of PD1^+^CD8^+^Glypican^+^CD103^+^ hybrid cells present in each tumour sample.

There is a positive association between the tumour size and overall survival time and between hybrid cell proportion and overall survival time (i.e. there is increased risk of death for higher tumour size and for higher hybrid cell proportions) (Table 4). We may infer with 95% confidence that the death rate from High hybrid cell proportion is approximately 2 times, and at least 1.13 times, the risk from Low hybrid cell proportion, holding tumour size constant (Table 5). Similarly, there is an 8% increase in the expected hazard relative to a one cm increase in tumour size, holding hybrid cell proportion constant (Table 5). By taking four risk factors including tumour stage, hybrid cell proportion, tumour size and patient age together, the multiple regression showed that the hybrid cell proportion and tumour size are still significant predictors of overall survival time, independently of others clinical risk factors (Table 4).

**Table 4:**
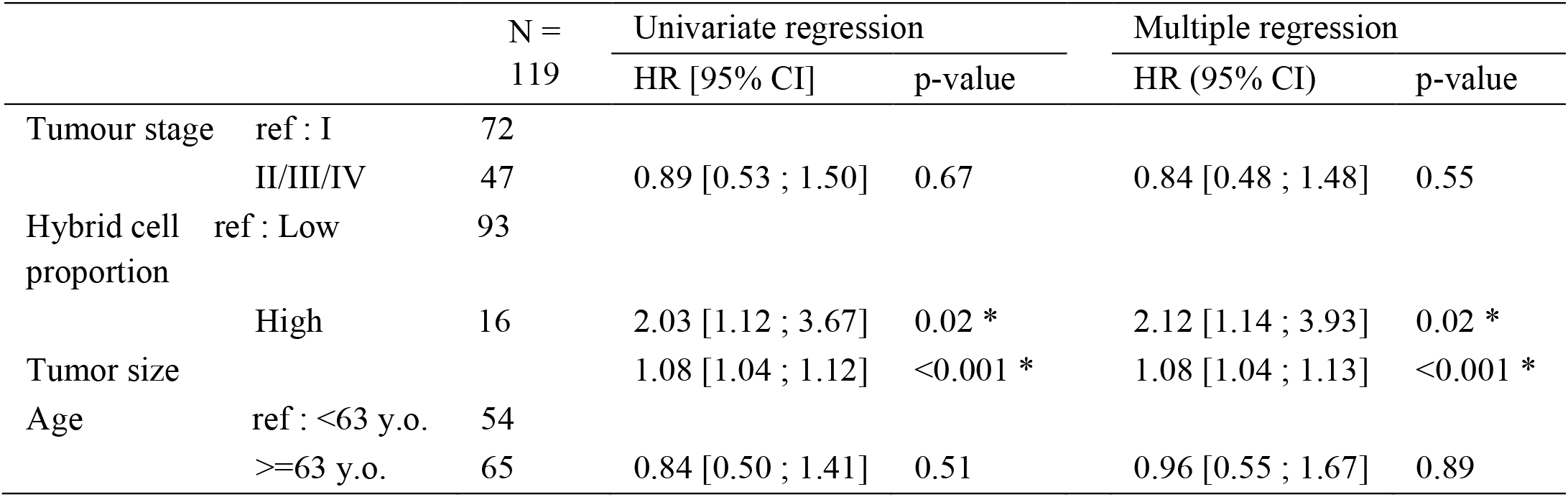
Univariate regression and multiple regression with tumour stage, hybrid cell proportion, tumour size and patient age as predictors of overall survival.

**Table 5:**
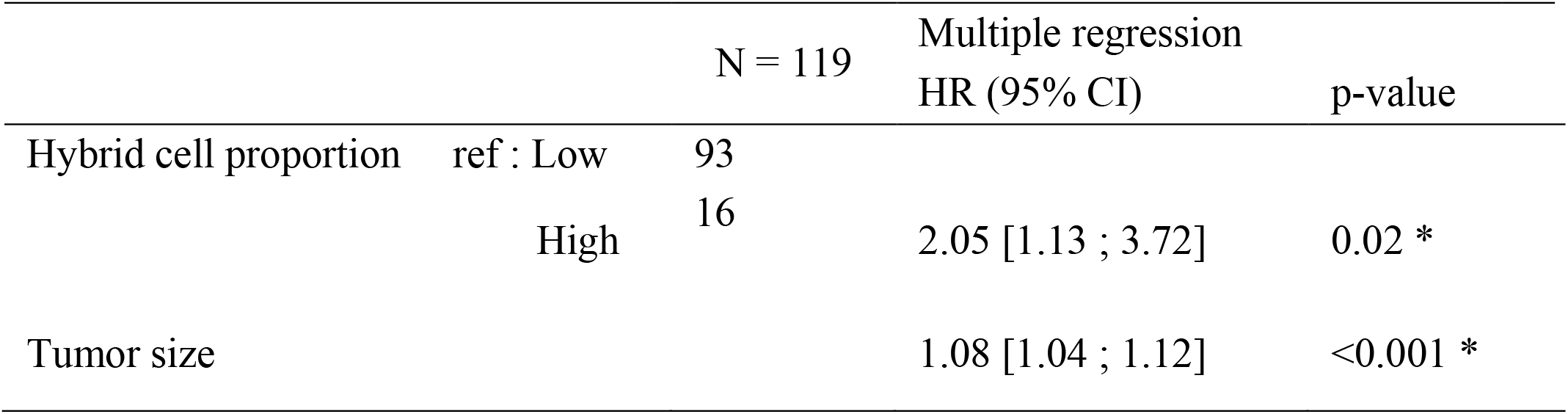
Multiple regression with hybrid cell proportion and tumour size as predictors of overall survival

### “Virtual reality” and “Augmented reality”

To further study the tumour-immune hybrid phenomenon, we developed our in-house innovative spatial lineage analytic pipeline known as “Virtual Reality (VR)”, which is also integrated into “Harmony”. Here, we incorporated the physical position of each cell and its corresponding FlowSOM unsupervised clustering-derived cell type into a XY cartesian plot (Patient A: Figure 11a and Patient B: Figure 11b). We combined the VR tool with the actual Vectra images to form “Augmented Reality (AR)”, where pathologists and immunologists can better identify the cell types based on their location (Figure 12). The resulting plots show the actual locations of the cells and their corresponding cell type, revealing important biological information about the tumour’s nature, such as whether there is immune-cell infiltration into the tumour sample. More importantly, these plots allow us to analyse the cellular distances between various cell types, which may hold important prognostic information.

**Figure 11:**
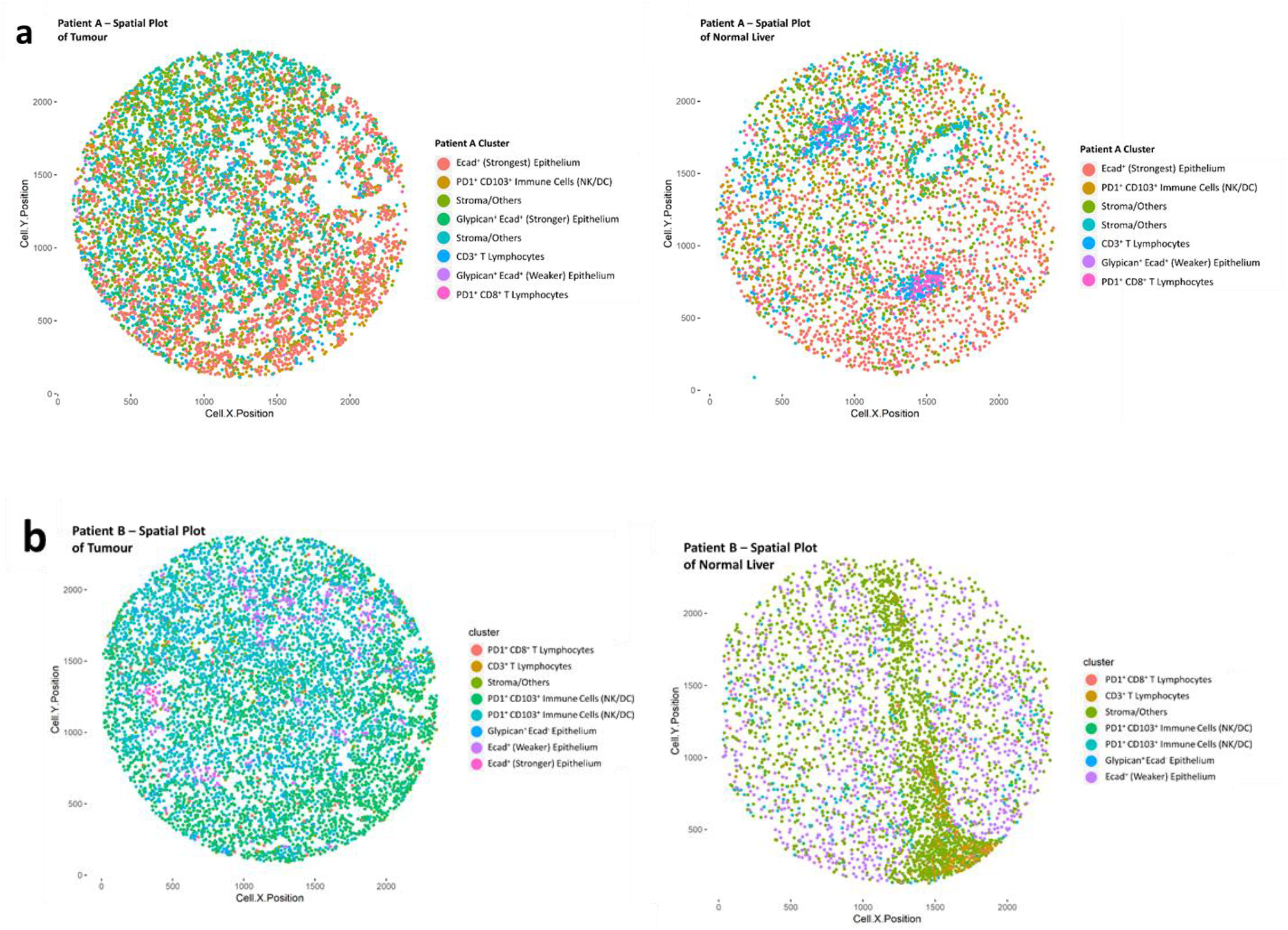
“Virtual Reality” – physical location of every cell with information regarding cell types displayed for a) Patient A; b) Patient B

**Figure 12:**
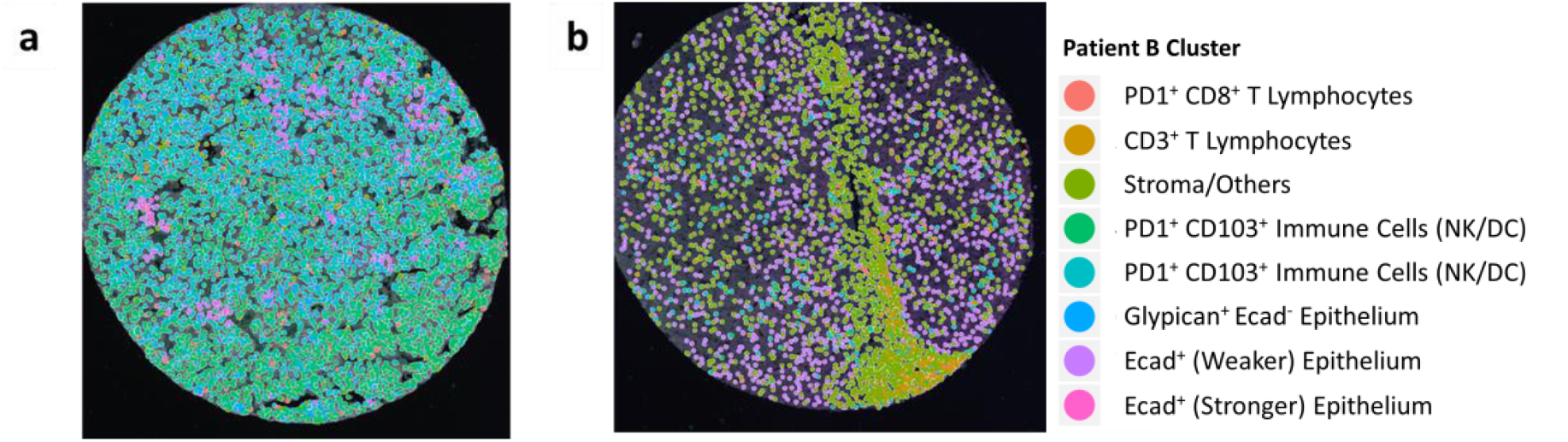
“Augmented Reality” – Patient B’s a) tumour and b) normal liver tissue’s physical location of every cell with information regarding cell types displayed and superimposed on Vectra images.

## Discussion

t-SNE and UMAP are both dimensionality reduction tools that have drawn considerable interest from the scientific community. Here, we show that both techniques have comparable performance, robustness and reproducibility, but UMAP has a clear advantage over t-SNE in terms of its runtime and preservation of biological lineages.

Consistent with previous findings, we found that the UMAP runtime is faster than t-SNE by approximately an order of magnitude (Figure 4a and 4b) ^27^. While the difference in timing may not be significant when handling small datasets, the difference is drastic when applied to datasets with a large number of cells or a large number of parameters, such as those derived from imaging mass cytometry or single-cell RNA sequencing. We also noted that the recently developed FIt-SNE does not improve the t-SNE visualisation runtime; however, as our dataset had a relatively low number of dimensions, we were unable to validate whether FIt-SNE performs better with a larger number of parameters. Interestingly, FIt-SNE and t-SNE had lower runtimes when running on the dataset with secondary features (total 112 parameters) compared to when running on primary features (seven parameters).

While others have reported that UMAP and t-SNE produce different clustering characteristics and shapes^6^, we instead observed similar clustering characteristics between both UMAP and t-SNE, with both methods giving rise to sufficiently distinct clusters (Figure 5). We hypothesize that this similarity is due to the lower number of biomarkers (only 7) adopted in our study. Variations between the two dimensionality reduction techniques may become more apparent when handling a higher number of biomarkers, leading to more diverse visualisations.

Moreover, one recent study claimed that the relative locations of clusters in UMAP represent the biological lineage between the cells^6^. However, from our study, we showed that the relative locations of the clusters are arbitrary and not fixed, like t-SNE (Figure 8). Hence, we believe that the physical locations of the clusters do not represent biological lineage, rather, the physical interactions between clusters, such as the bridging cluster 9 and arrangements of clusters provide information regarding the biological lineage (Figure 5d).

Interestingly, when we included the secondary features (with the same 7 biomarkers) from our study in the dimensionality reduction process, the t-SNE and UMAP plots showed virtually the same morphology (Figure 7). This concordance is likely due to greater clustering precision when more information regarding individual biomarker intensities in each cell is presented to the algorithm. This effect is currently unexplored in the field as the measurement features such as maximum, minimum and standard deviation of the marker intensities in the cell compartments are deemed less important than those that are currently widely used in the field. Mathematical parameters, such as standard deviation of the marker intensity, may reflect a cell’s immunologic nature. More research should be conducted in this area to determine whether this information is informative regarding the tumour immune microenvironment.

When using FlowSOM, we found strong Ecad expression and relatively weaker T-cell CD3 and CD103 marker expression in cluster number 8 from Patient B (Figure 3b). Despite some immune-cell labelling, however, these cells were still categorized as “Ecad^+^ (stronger) epithelium” as it is more likely that these cells are epithelial cells rather than immune cells. Vectra IF measures marker expression across a spectrum of intensities, making it hard to pinpoint the mean intensity cut-off to determine whether a marker is sufficiently present on the cells in the cluster. In other words, while the unsupervised cluster analysis may indicate the presence of a particular marker in that cluster, the values may be negligible in actual measurements. Moreover, there is the possibility of background noise that is induced in Vectra IF measurements due to the auto-fluorescent nature of liver cells.

We also noted from the UMAP and t-SNE visualisations of Patient B cluster 9 an interaction between Ecad^+^ epithelial cells and PD1^+^ CD8^+^ T lymphocytes (Figure 5c and 5d). We propose that this interaction is either due to immune cells losing immune markers, or tumour cells gaining immune markers, resulting in the presence of such a branch forming in the UMAP visualisation. However, both visualisation techniques were still capable of showing the presence of PD1^+^ CD8^+^ T lymphocytes on cluster 6. Recent data suggest that tumour cells express immune markers to evade detection by the body’s immune system ^28^. Having investigated this phenomenon by plotting the heatmaps of the biomarkers in this patient, we can see that the biomarker composition of these “bridging” cells are very likely tumour cells (Glypican^+^) with surface immune markers (PD1^+^, CD8^+^ and CD103^+^) (Figure 9). This points towards the possibility of tumour cells hybridising with CD8 T cells. The Kaplan-Meyer survival curve analysis also showed that patients with high proportions of these cells have poorer prognosis, supporting the study done by Gast et al (Figure 10). Moreover, multiple regression analysis shows that hybrid cell proportion is a significant factor associated with patient prognosis (Table 4 and 5). With this in mind, UMAP would be considered as superior to t-SNE in identifying cell lineages and may prove to be an important diagnostic feature for pathologists and immunologists to record when studying tumour sample characteristics.

With the advent of the cancer hybrid cell theory, new innovative ways to identify hybrid tumour cells from a biopsy sample would be of importance. Most excitingly, our new “VR” system can be used to complement traditional IHC to improve cell-type identification accuracy (Figure 12). These physical features, when complemented with the right markers may help determine whether tumour tissue is inflamed with myeloid or lymphoid cells, or non-inflamed by looking at the presence and location of immune cells. Integrating tumour-cell spatial and marker-based information will help drive research on personalised cancer treatments. Application of anti-glypican 3/CD3 bispecific T-cell-redirecting antibody treatment ^29^ is one example of a personalised treatment that may benefit from our VR approach. Identifying Glypican^+^ cells and T cells with their corresponding locations will allow better analysis on the feasibility of the therapy by tracking the cellular positions and determining whether there is a physical interaction between the two entities. Another possible application is in terms of PD1 marker expression. PD1 is a promiscuous marker that identifies macrophages, immune cells and tumour cells, making accurate cell-type identification difficult. Combining the VR tool with the actual Vectra images to form AR allows pathologists and immunologists to instead identify the cell types based on their location (Figure 12). Using the physical location of the cell, we can also study the prognostic characteristic of distances between tumour cells and key immune cells. VR thus has the potential to be employed as a new diagnostic tool to study tumour characteristics in the future.

## Conclusion

In summary, we believe that UMAP and t-SNE have very similar clustering capabilities when applied on Vectra IF data with 7 biomarkers. However, UMAP has a clear advantage over t-SNE in terms of its runtime. Moreover, there is a likelihood that UMAP can display cell biological lineages better than t-SNE given its more continuous plots as compared to t-SNE. We thus propose that UMAP can be used as an alternative to t-SNE. Adding on, we have also identified potential hybrid tumour cells that may contribute to worse prognosis among hepatocellular carcinoma patients. We have also developed VR and AR, which are spatial visualisation tools that allow pathologists and immunologists to identify the exact location of all the cell types. We have combined FlowSOM, UMAP, t-SNE, VR and AR as an automated R package analytical pipeline called “Harmony”. We strongly believe that “Harmony” has the potential to provide Vectra users with a qualitative analysis that can be complemented with Vectra IF’s quantitative results to provide pathologists and immunologists with more arsenal of tools to analyse tumour samples.

## Data availability

Raw input data and “Harmony” R package are available at https://github.com/JinmiaoChenLab/Harmony

## Acknowledgements

Grant information:

This study is supported by Singapore Immunology Network (SIgN) core fund.

## Author Contributions

Will change the names into the proper format later on:

Joe and Jinmiao conceived the study. Joe, Jinmiao and Duoduo designed the study methodology. Joe and Josh conducted the mIF and collected all patient HCC data. Jinmiao and Duoduo created “Harmony” with assistance from Grace. Duoduo performed all data analysis using “Harmony” except for multiple regression analysis, which was conducted by Marion. Joe and Jinmiao analysed the results. The manuscript was drafted by Duoduo and Joe, with key revisions inputted by Jinmiao.

## Supplementary Information

**Supplementary Figure 1:**
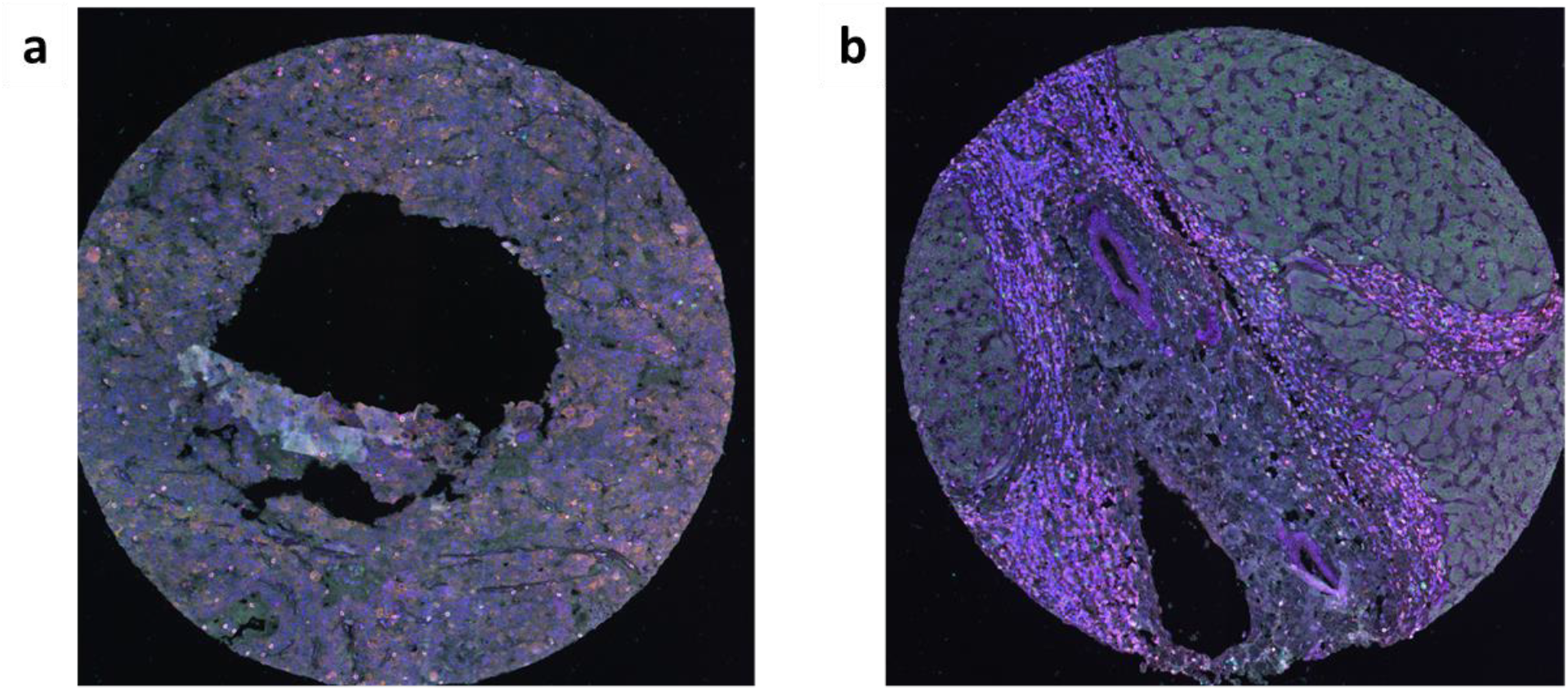
Additional Vectra images taken for patient B. a) Tissue sample taken from patient B’s tumour. b) Tissue sample taken from patient B’s region of normal liver.

**Supplementary Figure 2:**
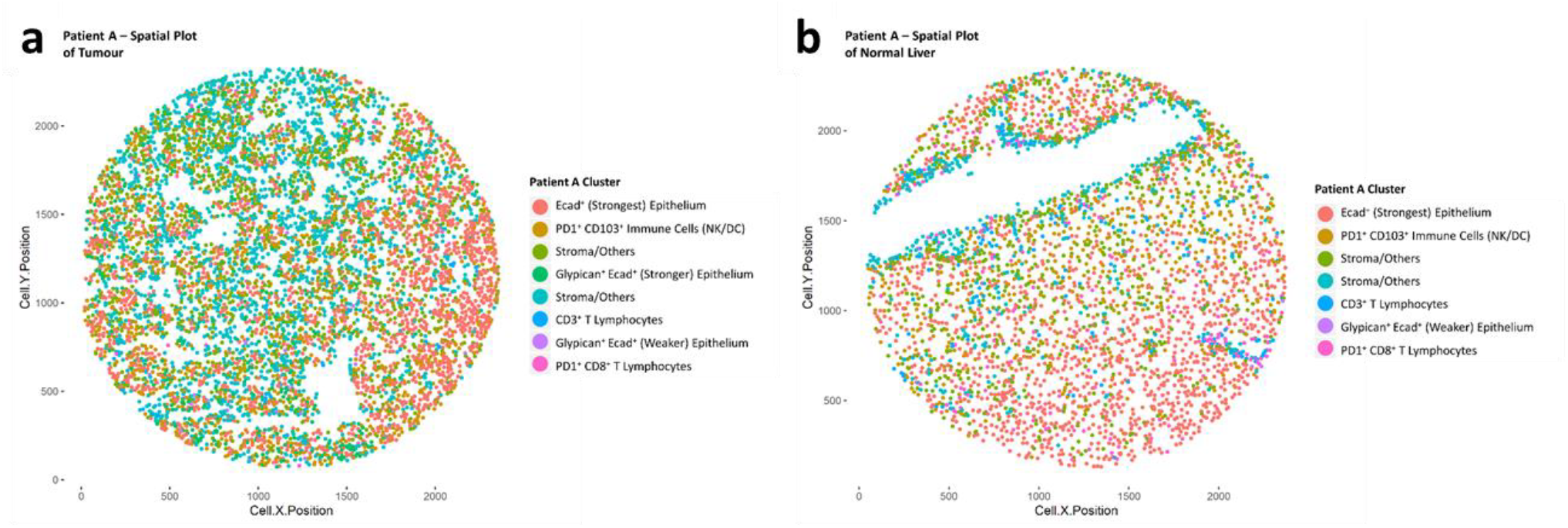
Patient A’s physical location of every cell with information regarding cell types displayed for additional Vectra images taken – “Virtual Reality” from 2 locations on the tumour (a) and normal liver (b).

**Supplementary Figure 3:**
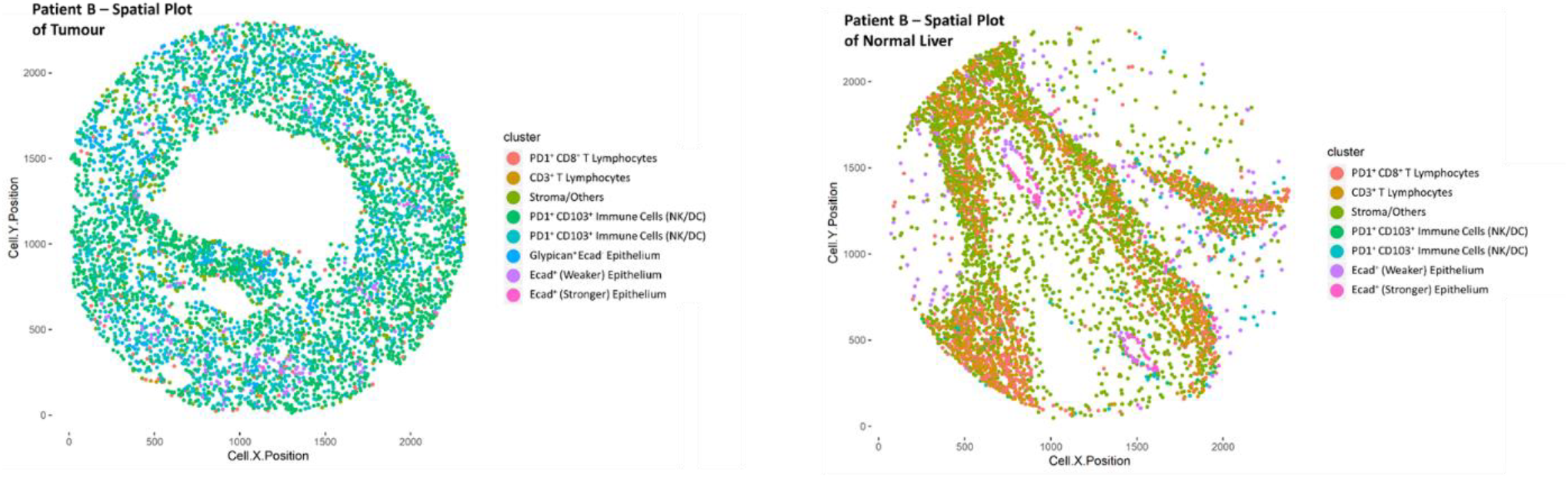
Patient B’s physical location of every cell with information regarding cell types displayed for additional Vectra images taken – “Virtual Reality”

**Supplementary Table 1:** All parameters measured using Vectra IHC

| Name of Feature Measured through Multiplex-Immunofluorescence |  |  |
| --- | --- | --- |
| **Primary Features**<br>**(7 in total)** | Mean intensity of cytoplasm CD3 | Mean intensity of cytoplasm PD1 |
|  | Mean intensity of cytoplasm CD8 | Mean intensity of cytoplasm CD103 |
|  | Mean intensity of cell membrane Ecad | Mean intensity of nucleus DAPI |
|  | Mean intensity of cytoplasm Glypican |  |
| **Secondary Features**<br>**(105 in total)** | Maximum intensity of cytoplasm CD3 | Maximum intensity of nucleus CD103 |
|  | Minimum intensity of cytoplasm CD3 | Minimum intensity of nucleus CD103 |
|  | Standard deviation of cytoplasm CD3 | Standard deviation of nucleus CD103 |
|  | Mean intensity of cell membrane CD3 | Mean intensity of entire cell CD103 |
|  | Maximum intensity of cell membrane CD3 | Maximum intensity of entire cell CD103 |
|  | Minimum intensity of cell membrane CD3 | Minimum intensity of entire cell CD103 |
|  | Standard deviation of cell membrane CD3 | Standard deviation of entire cell CD103 |
|  | Mean intensity of nucleus CD3 | Mean intensity of cytoplasm Ecad |
|  | Maximum intensity of nucleus CD3 | Maximum intensity of cytoplasm Ecad |
|  | Minimum intensity of nucleus CD3 | Minimum intensity of cytoplasm Ecad |
|  | Standard deviation of nucleus CD3 | Standard deviation of cytoplasm Ecad |
|  | Mean intensity of entire cell CD3 | Maximum intensity of cell membrane Ecad |
|  | Maximum intensity of entire cell CD3 | Minimum intensity of cell membrane Ecad |
|  | Minimum intensity of entire cell CD3 | Standard deviation of cell membrane Ecad |
|  | Standard deviation of entire cell CD3 | Mean intensity of nucleus Ecad |
|  | Maximum intensity of cytoplasm CD8 | Maximum intensity of nucleus Ecad |
|  | Minimum intensity of cytoplasm CD8 | Minimum intensity of nucleus Ecad |
|  | Standard deviation of cytoplasm CD8 | Standard deviation of nucleus Ecad |
|  | Mean intensity of cell membrane CD8 | Mean intensity of entire cell Ecad |
|  | Maximum intensity of cell membrane CD8 | Maximum intensity of entire cell Ecad |
|  | Minimum intensity of cell membrane CD8 | Minimum intensity of entire cell Ecad |
|  | Standard deviation of cell membrane CD8 | Standard deviation of entire cell Ecad |
|  | Mean intensity of nucleus CD8 | Maximum intensity of cytoplasm Glypican |
|  | Maximum intensity of nucleus CD8 | Minimum intensity of cytoplasm Glypican |
|  | Minimum intensity of nucleus CD8 | Standard deviation of cytoplasm Glypican |
|  | Standard deviation of nucleus CD8 | Mean intensity of cell membrane Glypican |
|  | Mean intensity of entire cell CD8 | Maximum intensity of cell membrane Glypican |
|  | Maximum intensity of entire cell CD8 | Minimum intensity of cell membrane Glypican |
|  | Minimum intensity of entire cell CD8 | Standard deviation of cell membrane Glypican |
|  | Standard deviation of entire cell CD8 | Mean intensity of nucleus Glypican |
|  | Maximum intensity of cytoplasm PD1 | Maximum intensity of nucleus Glypican |
|  | Minimum intensity of cytoplasm PD1 | Minimum intensity of nucleus Glypican |
|  | Standard deviation of cytoplasm PD1 | Standard deviation of nucleus Glypican |
|  | Mean intensity of cell membrane PD1 | Mean intensity of entire cell Glypican |
|  | Maximum intensity of cell membrane PD1 | Maximum intensity of entire cell Glypican |
|  | Minimum intensity of cell membrane PD1 | Minimum intensity of entire cell Glypican |
|  | Standard deviation of cell membrane PD1 | Standard deviation of entire cell Glypican |
|  | Mean intensity of nucleus PD1 | Mean intensity of cytoplasm DAPI |
|  | Maximum intensity of nucleus PD1 | Maximum intensity of cytoplasm DAPI |
|  | Minimum intensity of nucleus PD1 | Minimum intensity of cytoplasm DAPI |
|  | Standard deviation of nucleus PD1 | Standard deviation of cytoplasm DAPI |
|  | Mean intensity of entire cell PD1 | Mean intensity of cell membrane DAPI |
|  | Maximum intensity of entire cell PD1 | Maximum intensity of cell membrane DAPI |
|  | Minimum intensity of entire cell PD1 | Minimum intensity of cell membrane DAPI |
|  | Standard deviation of entire cell PD1 | Standard deviation of cell membrane DAPI |
|  | Maximum intensity of cytoplasm CD103 | Maximum intensity of nucleus DAPI |
|  | Minimum intensity of cytoplasm CD103 | Minimum intensity of nucleus DAPI |
|  | Standard deviation of cytoplasm CD103 | Standard deviation of nucleus DAPI |
|  | Mean intensity of cell membrane CD103 | Mean intensity of entire cell DAPI |
|  | Maximum intensity of cell membrane CD103 | Maximum intensity of entire cell DAPI |
|  | Minimum intensity of cell membrane CD103 | Minimum intensity of entire cell DAPI |
|  | Standard deviation of cell membrane CD103 | Standard deviation of entire cell DAPI |
|  | Mean intensity of nucleus CD103 |  |

## References

1. Lu, Y. et al. Dynamics of helper CD4 T cells during acute and stable allergic asthma. Mucosal Immunology (2018).

2. Tenenbaum, J.B., De Silva, V. & Langford, J.C. A global geometric framework for nonlinear dimensionality reduction. science 290, 2319–2323 (2000).

3. Coifman, R.R. et al. Geometric diffusions as a tool for harmonic analysis and structure definition of data: Diffusion maps. Proceedings of the National Academy of Sciences of the United States of America 102, 7426–7431 (2005).

4. Maaten, L.v.d. & Hinton, G. Visualizing data using t-SNE. Journal of machine learning research 9, 2579–2605 (2008).

5. McInnes, L. & Healy, J. Umap: Uniform manifold approximation and projection for dimension reduction. arXiv preprint arXiv:1802.03426 (2018).

6. Becht, E. et al. Dimensionality reduction for visualizing single-cell data using umap. Nature biotechnology 37, 38 (2019).

7. Van Gassen, S. et al. FlowSOM: Using self-organizing maps for visualization and interpretation of cytometry data. Cytometry. Part A: the journal of the International Society for Analytical Cytology 87, 636–645 (2015).

8. Lakhani, S., Ellis, I., Schnitt, S., Tan, P. & Van de Vijver, M. World Health Organisation Classification of Tumors of the Breast. International Agency for Research on Cancer 4, 142–147 (2012).

9. Ishak, K. et al. Histological grading and staging of chronic hepatitis. Journal of hepatology 22, 696–699 (1995).

10. Goodman, Z.D. Grading and staging systems for inflammation and fibrosis in chronic liver diseases. Journal of hepatology 47, 598–607 (2007).

11. Thike, A.A. et al. Loss of androgen receptor expression predicts early recurrence in triple-negative and basal-like breast cancer. Modern pathology: an official journal of the United States and Canadian Academy of Pathology, Inc 27, 352–360 (2014).

12. Stack, E.C., Wang, C., Roman, K.A. & Hoyt, C.C. Multiplexed immunohistochemistry, imaging, and quantitation: a review, with an assessment of Tyramide signal amplification, multispectral imaging and multiplex analysis. Methods 70, 46–58 (2014).

13. Abel, E.J. et al. Analysis and validation of tissue biomarkers for renal cell carcinoma using automated high-throughput evaluation of protein expression. Human pathology 45, 1092–1099 (2014).

14. Lovisa, S. et al. Epithelial-to-mesenchymal transition induces cell cycle arrest and parenchymal damage in renal fibrosis. Nature medicine 21, 998–1009 (2015).

15. Garnelo, M. et al. Interaction between tumour-infiltrating B cells and T cells controls the progression of hepatocellular carcinoma. Gut 15, 2015–310814 (2015).

16. Yeong, J. et al. Higher densities of Foxp3(+) regulatory T cells are associated with better prognosis in triple-negative breast cancer. Breast Cancer Res. Treat. 163, 21–35 (2017).

17. Garnelo, M. et al. Interaction between tumour-infiltrating B cells and T cells controls the progression of hepatocellular carcinoma. Gut 66, 342–351 (2017).

18. Lim, J.C.T., Yeong, J. P. S., Lim, C. J., Ong, C. C. H., Chew, V. S. P., Ahmed, S. S., Tan, P. H., & Iqbal, J. An automated staining protocol for 7-colour immunofluorescence of human tissue sections for diagnostic and prognostic use. Journal of The Royal College of Pathologists of Australasia (In Press).

19. Esbona, K. et al. COX-2 modulates mammary tumor progression in response to collagen density. Breast Cancer Research 18, 35 (2016).

20. Mlecnik, B. et al. The tumor microenvironment and Immunoscore are critical determinants of dissemination to distant metastasis. Sci Transl Med 8 (2016).

21. Nghiem, P.T. et al. PD-1 Blockade with Pembrolizumab in Advanced Merkel-Cell Carcinoma. New Engl J Med 374, 2542–2552 (2016).

22. Feng, Z. et al. Multispectral Imaging of T and B Cells in Murine Spleen and Tumor. The Journal of Immunology 196, 3943–3950 (2016).

23. Fiore, C. et al. Utility of multispectral imaging in automated quantitative scoring of immunohistochemistry. J. Clin. Pathol. 65, 496–502 (2012).

24. Abel, E.J. et al. Analysis and validation of tissue biomarkers for renal cell carcinoma using automated high-throughput evaluation of protein expression. Hum. Pathol. 45, 1092–1099 (2014).

25. Feng, Z. et al. Multiparametric immune profiling in HPV–oral squamous cell cancer. JCI Insight 2 (2017).

26. Linderman, G.C., Rachh, M., Hoskins, J.G., Steinerberger, S. & Kluger, Y. Efficient Algorithms for t-distributed Stochastic Neighborhood Embedding. arXiv preprint arXiv:1712.09005 (2017).

27. Becht, E. et al. Dimensionality reduction for visualizing single-cell data using UMAP. Nature Biotechnology (2018).

28. Gast, C.E. et al. Cell fusion potentiates tumor heterogeneity and reveals circulating hybrid cells that correlate with stage and survival. Science advances 4, eaat7828 (2018).

29. Ishiguro, T. et al. An anti–glypican 3/CD3 bispecific T cell–redirecting antibody for treatment of solid tumors. Science Translational Medicine 9 (2017).

